# Neurophysiological evidence of sensory prediction errors driving speech sensorimotor adaptation

**DOI:** 10.1101/2023.10.22.563504

**Authors:** Kwang S. Kim, Leighton B. Hinkley, Kurtis Brent, Jessica L. Gaines, Alvincé L. Pongos, Saloni Gupta, Corby L. Dale, Srikantan S. Nagarajan, John F. Houde

## Abstract

The human sensorimotor system has a remarkable ability to quickly and efficiently learn movements from sensory experience. A prominent example is sensorimotor adaptation, learning that characterizes the sensorimotor system’s response to persistent sensory errors by adjusting future movements to compensate for those errors. Despite being essential for maintaining and fine-tuning motor control, mechanisms underlying sensorimotor adaptation remain unclear. A component of sensorimotor adaptation is implicit (i.e., the learner is unaware of the learning process) which has been suggested to result from sensory prediction errors–the discrepancies between predicted sensory consequences of motor commands and actual sensory feedback. However, to date no direct neurophysiological evidence that sensory prediction errors drive adaptation has been demonstrated. Here, we examined prediction errors via magnetoencephalography (MEG) imaging of the auditory cortex (n = 34) during sensorimotor adaptation of speech to altered auditory feedback, an entirely implicit adaptation task. Specifically, we measured how speaking-induced suppression (SIS)--a neural representation of auditory prediction errors--changed over the trials of the adaptation experiment. SIS refers to the suppression of auditory cortical response to speech onset (in particular, the M100 response) to self-produced speech when compared to the response to passive listening to identical playback of that speech. SIS was reduced (reflecting larger prediction errors) during the early learning phase compared to the initial unaltered feedback phase. Furthermore, reduction in SIS positively correlated with behavioral adaptation extents, suggesting that larger prediction errors were associated with more learning. In contrast, such a reduction in SIS was not found in a control experiment in which participants heard unaltered feedback and thus did not adapt. In addition, in some participants who reached a plateau in the late learning phase, SIS increased (reflecting smaller prediction errors), demonstrating that prediction errors were minimal when there was no further adaptation. Together, these findings provide the first neurophysiological evidence for the hypothesis that prediction errors drive human sensorimotor adaptation.

## Introduction

The sensorimotor system shows a remarkable ability to quickly and efficiently learn movements based on sensory feedback. Soon after perceiving sensory errors that arise from movements, the system updates future movements to compensate for the errors, a phenomenon called sensorimotor adaptation. What drives such an elegant learning process? Previous studies suggested that adaptation can be driven by both task errors (i.e., discrepancy between the action and the goal) and sensory prediction errors (i.e., mismatches between the actual sensory consequences of a movement and those predicted from the motor commands driving that movement).

In the speech domain, however, multiple lines of evidence suggest that speech sensorimotor adaptation to altered auditory feedback is implicit (i.e., participants are unaware of the learning), which is thought to be driven mainly by sensory prediction errors (Mazzoni & Krakauer, 2006). In previous adaptation studies, participants showed no difference in the amount of learning in response to formant-perturbed auditory feedback when instructed to compensate, to ignore the feedback, or to avoid compensating (Keough et al., 2013; Munhall et al., 2009). Although behavioral studies have suggested that this unconscious minimizing of auditory prediction errors is the signal that drives speech sensorimotor adaptation, direct neurophysiological evidence of this process has not been demonstrated.

One neural representation of auditory prediction errors is speaking-induced suppression (SIS) of the auditory cortex. Studies have reported that the auditory responses to self-produced speech are smaller (i.e., suppressed) than the responses to playback of the same speech sound, consistent with the idea that auditory responses arise from auditory prediction errors, which are small in the self-produced case (i.e., auditory feedback is predictable) and large in the passively heard case (i.e., auditory feedback is unpredictable). Thus, SIS demonstrates that, during speaking, the auditory system predicts and anticipates the arrival of auditory feedback of speech onset, resulting in a suppressed feedback comparison response, as compared to auditory responses during passive listening to playback when speech onset cannot be predicted/anticipated. Consistent with the idea, SIS was reduced when participants spoke with pitch-perturbed auditory feedback (e.g., Behroozmand & Larson, 2011; Chang et al., 2013) or voice-manipulated auditory feedback (“alien voice”, e.g., Heinks-Maldonado et al., 2005, 2006; Houde et al., 2002). Importantly, this reduction in the suppression of auditory areas in response to perturbed auditory feedback are not unique to human speech, as they have also been observed in marmoset monkey vocal production (e.g., Eliades & Tsunada, 2018).

Previously, reduction in a similar suppression effect (i.e., suppressed neural response in active movements compared to passive movements) has been found in Rhesus monkey cerebellum during sensorimotor adaptation (Brooks et al., 2015), but no such evidence has been documented in humans to date. One previous study that examined SIS during adaptation to first formant frequency shifts via electroencephalography (EEG) reported that SIS amplitude in the learning phase (i.e., during perturbed first formant) was not reduced compared to the pre-adaptation baseline (Sato & Shiller, 2018). However, the negative finding could result from masking of SIS changes across all 80 feedback perturbation trials, as opposed to changes that may have occurred in early trials (e.g., initial 20 to 40 feedback perturbation trials) when most adaptation occurs (e.g., Kim & Max, 2021). Here, we used magnetoencephalography (MEG) imaging during repeated speech adaptation sessions to test the hypotheses that (1) SIS reduces during early phases of speech sensorimotor adaptation, and (2) the early SIS reduction may be distinct from SIS changes found in later phases of adaptation.

## Results

Participants lay supine on the scanner bed of a whole-head, 275-channel biomagnetometer system (MEG; Omega 2000, CTF, Coquitlam, BC, Canada) for a total of four sessions (first and second speaking sessions, first and second listening sessions). During the first two sessions, participants were asked to read “Ed,” “end,” or “ebb” (60 trial blocks for 3 different words = 180 total trials) that appeared on the screen. During these speaking sessions, participants heard their speech with the first formant frequency (Formant 1 or F1) shifted upward for some trials, which made their speech to sound like “add,” “and,” and “abb,” respectively. Specifically, after the first 20 trial blocks (i.e., baseline) which had no perturbation, the 150 Hz up-shift perturbation was present from trial block 21 to 50. We categorized the first 15 trial blocks of the perturbed trials (21 – 45) as the early learning phase and the second 15 trial blocks (36-50) as the late learning phase.

After the first session, participants were given a few minute-long break that included conversations with the experimenter, which allowed additional exposure to their unaltered auditory feedback (Figure 1). We then asked participants to repeat another speaking session. The rationale for this repeated session was that most adaptation occurs quickly, often in the first 10-30 trials of the perturbation phase, but such a low number of trials does not provide enough power for the evoked potential analyses. Thus, to ensure an adequate number of trials for the early and late learning phases, an additional session was recorded. After completing two speaking sessions, participants were asked to listen to their recorded speech in the first two speaking sessions across the subsequent two sessions (i.e., listening sessions). During the listening sessions, participants saw the same stimuli (i.e., words) that they saw in the speaking sessions (see Methods for more details).

**Figure 1.**
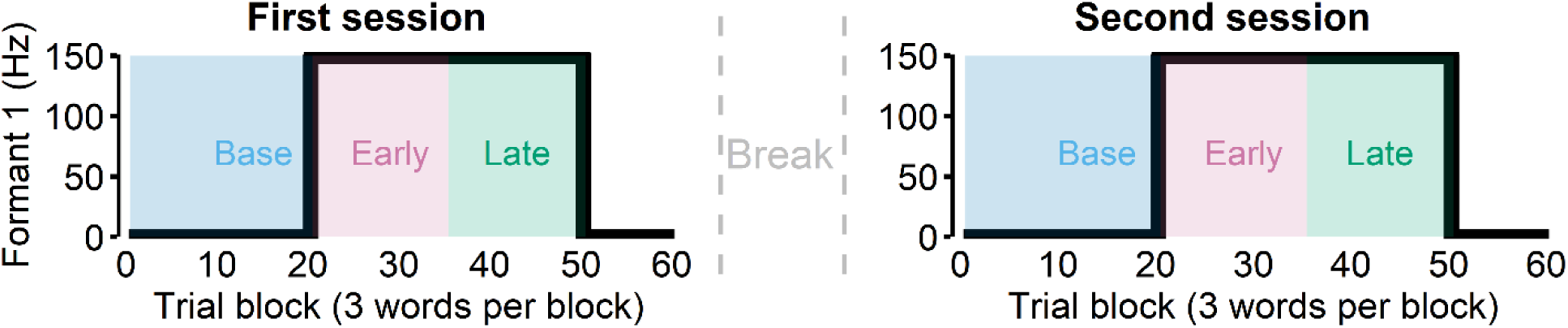
Participants were asked to read words during the first two sessions (“speak”). In these sessions, 150 Hz up-shift perturbation was present from the trial block 21 to 50. We categorized the first 15 trial blocks of the perturbed trials (21 – 45) as the early learning phase and the second 15 trial blocks (36-50) as the late learning phase. After the first session, we asked participants to repeat another speaking session after a break lasting a few minutes.

We averaged the acoustic and MEG data across the repeated sessions. As shown in Figure 2, source localization of trial-averaged data for each condition (speak, listen) and phase (baseline, early learning, and late learning) was conducted to determine peak activity (M100) location within the auditory cortex. We then computed the M100 amplitude differences between the listen and speak sessions to determine SIS for each condition and phase (see Methods for more details).

**Figure 2.**
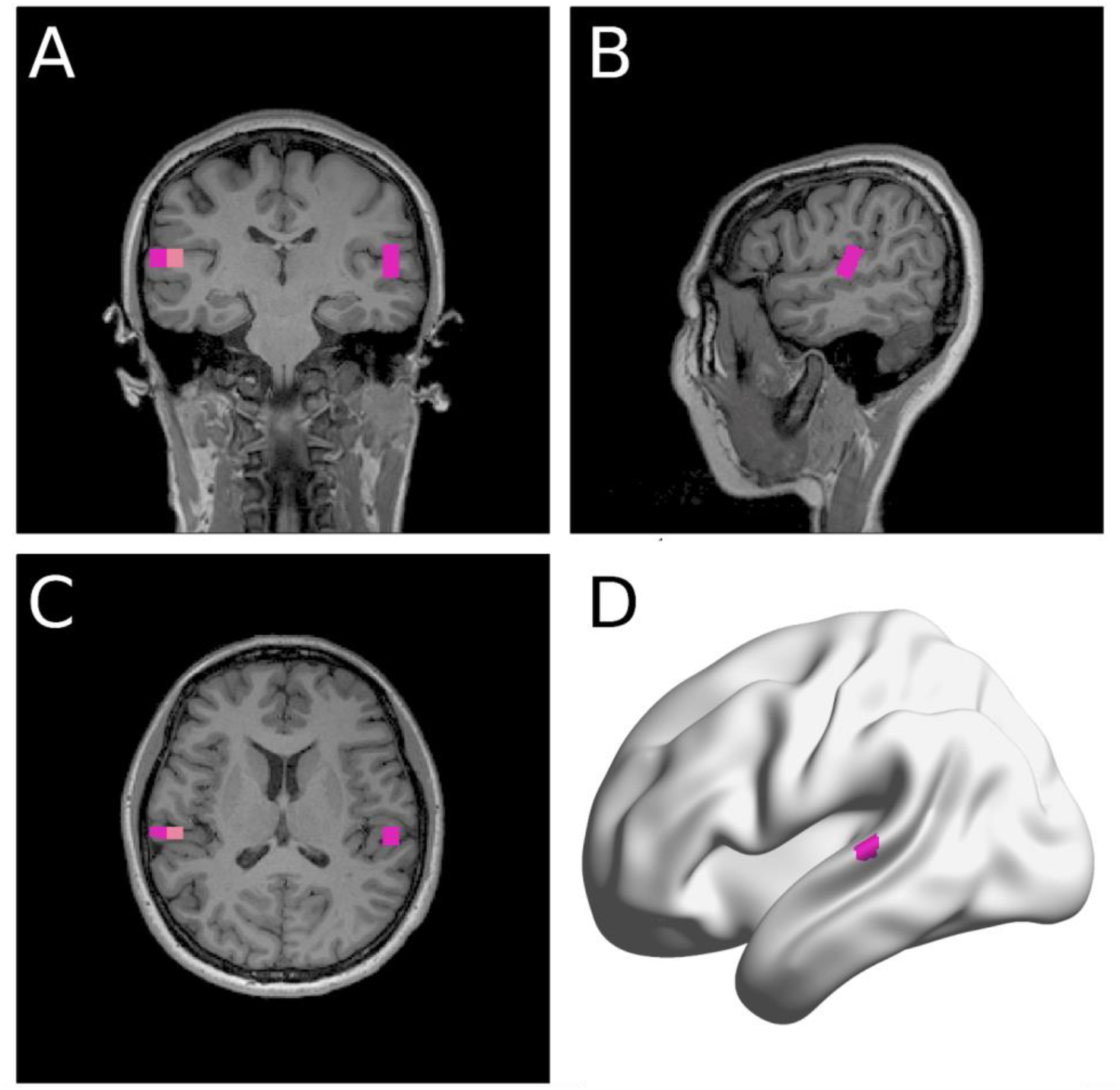
A representative participant’s source localization. NUTMEG (Hinkley et al., 2020) identified a few MNI coordinates that showed clear M100 response shown in the coronal (**A**), sagittal (**B**), and transverse (**C**) planes. The MNI coordinate of the voxel with the most power in the auditory areas in each hemisphere was selected for analyses. **D**: The same participant’s left auditory area coordinate selected shown in a surface-based rendering (BrainNet Viewer, Xia et al., 2013).

### SIS reduction was positively correlated with early adaptation

Nearly all participants adapted in both speaking sessions (Fig. 3A), except for three participants who adapted in only one of the two sessions. Given that there was no evidence of savings (i.e., changes in the baseline or learning behavior from repeating the task, see Supplementary Information 1), these participants were included in the analyses. The SIS analyses revealed that there was no right hemisphere SIS (see Supplementary Information 2), which is known to be variable across tasks and individuals (see Discussion for more details). On the other hand, most participants showed a clear suppression of left auditory activity in the speaking condition (compared to the listening condition) during the baseline phase (Fig. 3B, left). Hence, SIS refers to suppression of *left* auditory activity hereafter unless specified otherwise.

**Figure 3.**
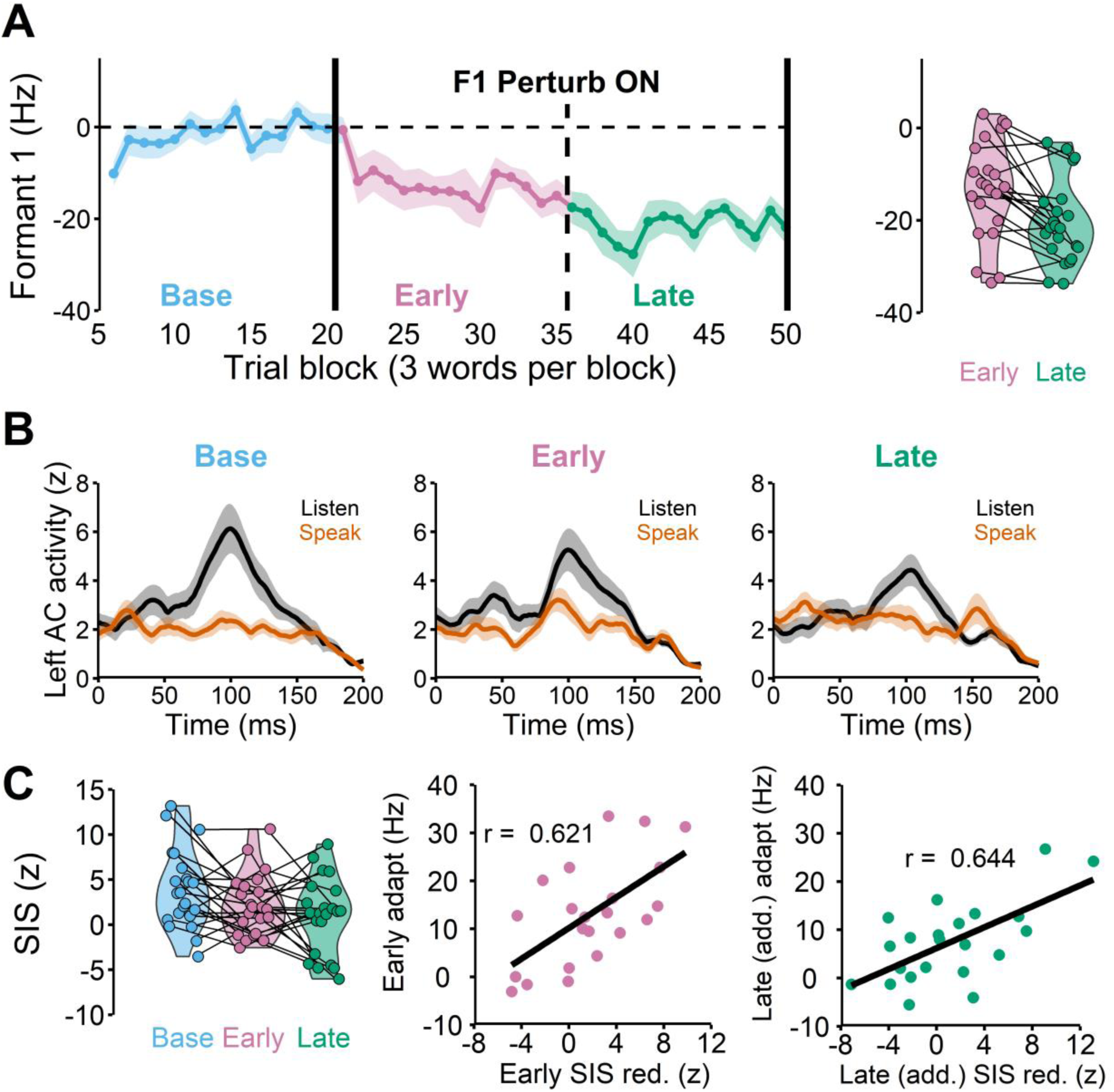
**A**: The group average speech auditory-motor adaptation in which participants lowered their first formant frequency (F1) in response to the 150 Hz upshift F1 perturbation (left). Each participant’s early and late adaptation. Some participants continued to adapt after the early phase, but others plateaued (right). **B**: The left auditory cortex responses (M100) in listen and speak conditions demonstrate that the amount of speaking-induced suppression (i.e., listen (black) – speak (orange)) is reduced during early learning (Early) compared to the baseline (Base). **C**: SIS was significantly reduced in the early and late learning phases compared to the baseline (left, r(20) = 0.621, p = 0.002). The amount of SIS reduction in the early learning phase was significantly correlated with the amount of early adaptation (middle). The amount of additional SIS reduction in the late learning phase also significantly correlated with the additional amount of adaptation in the phase (right, r(20) = 0.644, p = 0.001).

We also found that the SIS response changed across the baseline, early, and late learning phases (Fig. 3B, middle and right), F(2, 44) = 4.788, p = 0.013. The post-hoc pairwise comparison test indicated that SIS response in the early learning phase did not differ from the baseline (Fig. 3C, left), t(46.1) =1.829, p_adj_ = 0.171. Interestingly, we found that 14 participants who adapted more than 10 Hz significantly reduced SIS in the early learning phase, t(13) = – 2.903, p_adj_ = 0.025, whereas 8 participants who adapted less than 10 Hz showed no reduction in SIS, t(7) = –0.478, p_adj_ = 0.647. Indeed, the correlation coefficient between the amount of SIS reduction in the early learning phase and the amount of learning (in the early learning phase) across participants was significant, r(20) = 0.621, p = 0.002 (Fig. 3C, middle).

### Further SIS reduction was positively correlated with (additional) late learning

The SIS amplitude in the late learning phase was also significantly reduced compared to the baseline (Fig. 3C, left), t(46.1) = 4.339, p_adjust = 0.002. We found that the SIS reduction from the baseline was not significantly correlated, though trending, with the final amount of adaptation in the late learning phase, r(20) = –0.416, p = 0.054 (see Supplementary Information 3). This result was somewhat consistent with our hypothesis that most learning typically occurs in the early phase, and thus the late phase SIS reduction from baseline would not be able to capture most of the adaptation extent. Rather, late SIS reduction that accounts for early SIS changes (i.e., *additional* late SIS reduction from early SIS) is likely a predictor for late (additional) learning behaviors. Indeed, we found that additional SIS reduction in the late learning phase (i.e., late SIS relative to the early SIS) was significantly correlated with additional late adaptation, i.e., late adaptation relative to early adaptation r(20) = 0.644, p = 0.001.

Another related finding is that there were 9 participants whose late learning SIS was not reduced compared to early SIS, but rather increased in the late learning phase, t(8) =5.539, p = 0.001. This SIS increase resulted in a near-complete SIS recovery to the baseline level (i.e., the late learning SIS response did not differ from the baseline SIS response), t(8) = –0.804, p = 0.445. Importantly, these participants also did not show a significant amount of additional learning in this late learning phase, t(8) = –1.425, p = 0.192, even though adaptation remained largely incomplete in these participants (i.e., adapted 13.6% of the perturbation size). Consistent with this view, 13 participants whose late learning SIS was reduced compared to early SIS (t(12) = 3.869, p = 0.002) showed continual adaptation (i.e., late learning significantly different from early learning), t(12) = –4.627, p = 0.001.

Taken together, the relationship between additional SIS reduction and adaptation in the late learning phase also followed the same trend found in the early learning phase. That is, individuals who showed more reduction in SIS, also tended to show more learning, suggesting that larger adaptation was associated with larger prediction errors. In contrast, less learning or no learning behavior (e.g., reaching a plateau) was associated with smaller prediction errors (i.e., increases in SIS).

### SIS remained unchanged when there was no learning

To ensure that SIS reduction was related to learning behaviors, we designed a control experiment in which there was no auditory perturbation (and thus no learning was expected). Here, participants also completed two speaking and two listening sessions. Other than the absence of the perturbation, the experimental setup and the analyses methods were identical to the main experiment. We found that participants did not adapt (Fig 4A) and SIS reduction also did not occur (i.e., SIS amplitudes did not change across the phases), F(2, 24) = 0.211, p = 0.812. Therefore, SIS remained unchanged when there was no learning.

**Figure 4.**
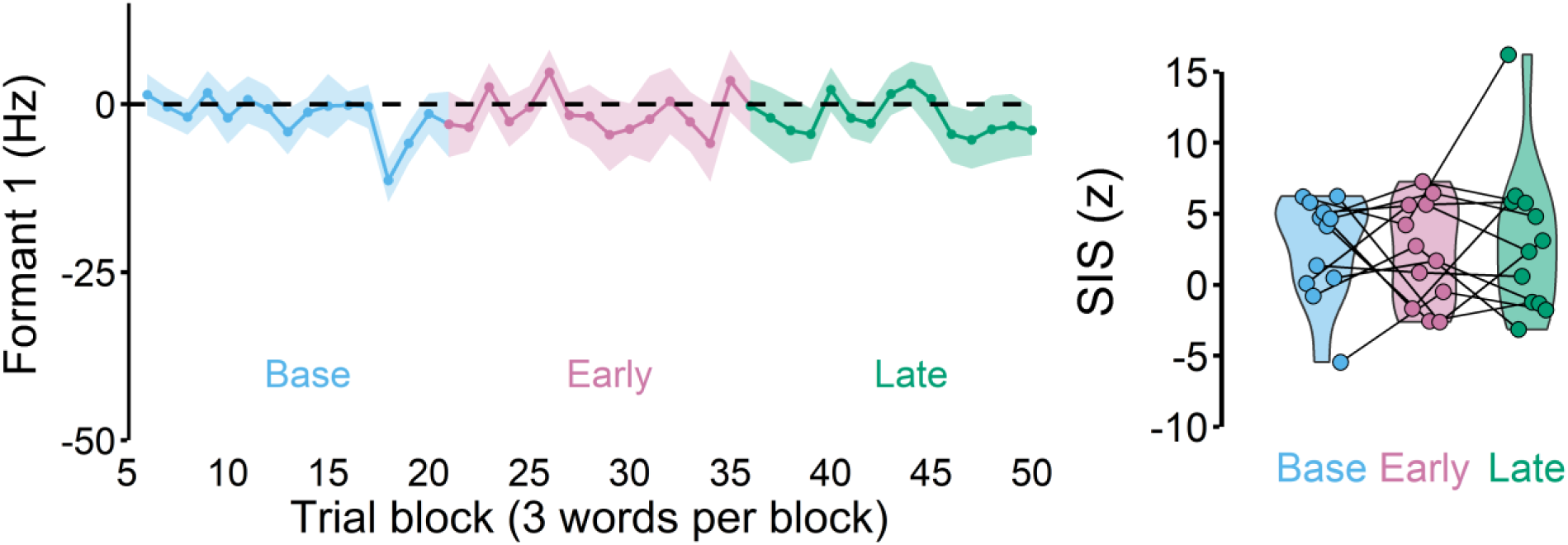
A control experiment in which no auditory perturbation was applied. As expected, participants did not show any changes in Formant 1, exhibiting, no learning (left). There was also no SIS change across the different phases (right).

## Discussion

We used magnetoencephalography (MEG) imaging to examine auditory prediction errors during speech auditory-motor adaptation. Specifically, we measured speaking-induced suppression (SIS)—suppression of auditory responses to self-produced speech compared to the responses to passively heard speech—which is thought to represent auditory prediction errors. To fully capture SIS changes in the early learning phase during which most of adaptation typically occurs, we analyzed the early learning and late learning phases separately.

### Neurophysiological evidence that auditory prediction errors drive implicit adaptation

SIS was significantly reduced in the early learning phase during which adaptation occurred. In contrast, in a control experiment in which there was no perturbation (and thus no adaptation), such a SIS reduction was not found. In addition, the amount of SIS reduction was positively correlated with the amount of adaptation, delineating a direct link between prediction errors (i.e., more SIS reduction) and adaptation. Furthermore, in the late learning phase, SIS increase, instead of SIS reduction, was associated with adaptation reaching an asymptote (i.e., absence of further learning). Hence, it is unlikely that SIS change arises simply because of adaptive behavior (i.e., lower F1 productions). Rather, SIS reduction likely reflects prediction error signals that lead to adaptation. In sum, our results suggest that auditory prediction errors *drive* speech auditory-motor adaptation.

Our findings are consistent with previous reports of speech adaptation being entirely implicit (e.g., Kim & Max, 2021; Lametti et al., 2020), which is thought to be driven by prediction errors (Haith & Krakauer, 2013; Mazzoni & Krakauer, 2006). In addition, speech adaptation also seems to be sensitive to auditory feedback delays (i.e., 100ms delay can eliminate adaptation), which highlights the importance of prediction errors that require temporally precise comparison of prediction and the actual feedback (Max & Maffett, 2015; Shiller et al., 2020). More recently, a computational model, Feedback-Aware Control of Tasks in Speech (FACTS, Parrell et al., 2019) also generated simulations of adaptation driven by auditory prediction errors (K. S. Kim et al., 2023). Recently, Tang et al. (2023) showed that SIS was changed after exposure to auditory perturbation that manipulated participants’ perceived variability. Given that this type of learning during which participants change production variability (Tang et al., 2022; Wang & Max, 2022) likely involves prediction errors as in the current study, the results of Tang et al. (2023) are consistent with our own findings.

To date, only one other study examined SIS during speech auditory-motor adaptation, but they reported no SIS changes during adaptation (Sato & Shiller, 2018). Although their finding may seem contradictory to the current study at first glance, it should be noted that in the previous study SIS amplitudes across the whole learning phase (80 trials) were averaged and analyzed together, which likely included SIS recovery response in the late phase as found in the current study’s late learning phase. Hence, it is possible that SIS reduction was present in the early learning phase, but such an effect may have been weakened by the late perturbation data.

It should be noted that our findings do not necessarily reject the notion that task errors may also drive implicit speech adaptation. In upper limb visuomotor rotation, recent studies have demonstrated that task errors contribute to implicit adaptation (Albert et al., 2022; H. E. Kim et al., 2019; Leow et al., 2018, 2020; Miyamoto et al., 2020; Morehead & Xivry, 2021). Although it remains possible that other types of errors (in addition to prediction errors) may also influence speech adaptation, such evidence has not been documented (also see “What does SIS reflect?” below).

Broadly, our findings provide the first neurophysiological evidence that sensory prediction errors drive implicit adaptation in humans. A similar suppression effect has been previously documented in the cerebellum of rhesus monkey during head movement adaptation (Brooks et al., 2015). In the study, cerebellar neuron activities, which are typically suppressed during voluntary movements compared to passive movements much like SIS, did not differ between the two conditions (voluntary vs. passive) during adaptation. Remarkably, this reduced suppression also recovered (i.e., suppression increased) towards later learning trials, directly in line with our result. Here, we expanded the previous finding by demonstrating that the extent of such suppression reduction (or recovery) was closely associated with implicit adaptation across individuals.

### Adaptation plateaus when prediction errors are minimal

Another interesting finding of the current study concerns a potential mechanism that causes adaptation to halt. In the past, several explanations for why adaptation is incomplete have been put forth, especially for speech adaptation which often plateaus around 20-40% (see Kitchen et al., 2022 for detailed discussion). Some studies have demonstrated that speech adaptation accompanies changes in perceptual boundaries which may contribute to incomplete adaptation (Lametti et al., 2014; Shiller et al., 2009), but perceptual auditory targets do not seem to change throughout adaptation (K. S. Kim & Max, 2021) and preventing perceptual target shifts by playing back the participants’ baseline productions did not increase adaptation. Others argued that a conflict between unperturbed somatosensory feedback and perturbed auditory feedback may lead to limited adaptation, but this account also lacks supporting evidence. In fact, preliminary data from our laboratory shows that even when somatosensory feedback becomes unreliable by oral application of lidocaine, adaptation behavior does not increase, suggesting that somatosensory feedback may not be a reason for incomplete adaptation.

One idea consistent with previous studies in upper limb reaching adaptation is that consistency of errors modulates error sensitivity, which results in limited adaptation (e.g., Albert et al., 2021). This idea has not been directly examined in the context of speech adaptation, but it is plausible that the overall size of prediction errors may be modulated by feedback (or perturbation) consistency. Some studies have found that individuals with high perceptual (auditory) acuity measured by psychometric functions had a larger extent of adaptation (e.g., Daliri & Dittman, 2019), which may suggest a potential link between error sensitivity and adaptation. However, several other studies failed to find such a relationship (e.g., Abur et al., 2018; Alemi et al., 2021; Feng et al., 2011; Lester-Smith et al., 2020).

Another potential explanation is that adaptation is halted by prediction errors which quickly decrease throughout adaptation because of both the motor output changes and sensory prediction updates, an idea put forth by a computational model, FACTS (K. S. Kim et al., 2023). In these simulations, the adaptive motor output produced lower F1 in response to F1 upshift perturbation, resulting in perturbed sensory feedback to become more like the baseline sensory feedback (i.e., lower perturbed feedback in F1). Interestingly, the simulations showed that sensory prediction was also updated to predict perturbed auditory feedback (i.e., higher prediction in F1). Thus, prediction errors, the difference between lower perturbed feedback in F1 and higher prediction in F1, became minimized throughout adaptation, eventually becoming a small amount that could no longer induce adaptation.

Empirical evidence for the idea that minimal prediction errors may result in halting adaptation can be found in head movement adaptation of rhesus monkeys (Brooks et al., 2015). In the study, cerebellar neuron activities to the voluntary head movement became more suppressed (compared to passive movement) as adaptation plateaued. Critically, the authors argued that the neural response becoming more suppressed (or less “sensitive”) throughout learning demonstrates that sensory prediction was being rapidly updated to predict unexpected (perturbed) sensory feedback.

In the current study, late learning phase SIS increased (i.e., minimal prediction errors) in multiple participants who also showed plateaued adaptation in the phase (i.e., no additional learning) which is directly in line with previous findings. Furthermore, the observation that adaptation plateaued even though adaptation was largely incomplete (i.e., 14.88% of the perturbation size) can be best explained by the idea that sensory forward model updates (i.e., prediction updates) may have occurred throughout adaptation, minimalizing prediction errors. Thus, our findings corroborate the notion that incomplete adaptation may result from not only the motor output changes but also sensory prediction updates, which together minimize prediction errors.

### What does SIS reflect?

SIS is typically viewed as a measure that reflects prediction errors given that SIS is reduced upon unexpected auditory feedback (e.g., pitch perturbation, alien voice, Heinks-Maldonado et al., 2005). This view is also shared by other studies examining suppression of motor-evoked auditory responses (i.e., finger pressing a button to generate a tone), which is also reduced or absent in deviant (i.e., unpredicted) sounds (Aliu et al., 2009; Knolle et al., 2013). In contrast to this view, a previous study from our laboratory argued that the SIS response may instead reflect target errors, discrepancies between an intended auditory target with auditory feedback (Niziolek et al., 2013). In the study, Niziolek and colleagues found that the greater the onset formants deviated from the median formants, the more SIS was reduced. Additionally, this reduction in SIS correlated with the amount of subsequent within-utterance formant change that reduced variance from the median as the utterance progressed (“centering”). Under the assumption that the median formants are close to the intended auditory target (i.e., an ideal production), it can be argued that SIS reflects target errors.

However, our finding that SIS increased in 9 participants during the late learning phase cannot be easily explained by this account. Due to the SIS recovery, their late learning phase SIS response, which did not differ from their baseline SIS response, would be interpreted as minimal or no target errors according to the target error explanation for SIS. Nonetheless, these participants compensated for only 13.6% of the perturbation on average, presumably leaving a considerable discrepancy between any fixed auditory target and auditory feedback. While previous studies have reported perceptual boundaries shifting towards the direction of perturbation during adaptation which may reduce target errors (Lametti et al., 2014; Shiller et al., 2009), it has also been suggested that auditory targets, as opposed to perceptual boundaries, do not change throughout adaptation (K. S. Kim & Max, 2021). In fact, a recent study has demonstrated that playing back the median production (i.e., the assumed auditory target) to participants throughout adaptation did not affect learning (LeBovidge et al., 2020), raising questions about whether auditory targets change during adaptation.

Alternatively, if SIS indeed reflects prediction errors rather than target errors, this view offers a different interpretation of Niziolek et al. (2013). According to the view, reduced SIS in productions with greater deviations from the median production may have resulted from large signal-dependent noise that stemmed from both the lower neural and muscular motor systems (Harris & Wolpert, 1998; Houde et al., 2014; Jones et al., 2002). Because such noise cannot be predicted by cortical areas, observed auditory feedback would not match auditory prediction, leading to large auditory prediction errors. Hence, it is plausible that the reduced SIS found in those productions reflects larger prediction errors. This view would also imply that centering (i.e., subsequent within-utterance formant change) minimized prediction errors, rather than target errors.

### Neural correlates of auditory prediction errors

In the current study, we estimated auditory prediction errors from activities in the auditory cortex. Given that auditory areas receive corollary discharge from speech motor areas during speech production (Khalilian-Gourtani et al., 2022), it is possible that auditory prediction errors may be computed in the auditory cortex. However, a large body of evidence suggests that the cerebellum may be a neural substrate for forward models that generate sensory predictions (e.g., Blakemore et al., 1999, 2001; Imamizu & Kawato, 2012; Kawato et al., 2003; Pasalar et al., 2006; Shadmehr, 2020; Shadmehr & Krakauer, 2008; Skipper & Lametti, 2021; Therrien & Bastian, 2019; Wolpert et al., 1998). Studies have also documented evidence that the cerebellum may also compute sensory prediction errors (e.g., Blakemore et al., 2001; Brooks et al., 2015; Cullen & Brooks, 2015). Alternatively, it has also been hypothesized that the cerebellum may work in concert with cortical areas to generate sensory prediction mechanisms and prediction errors (Blakemore & Sirigu, 2003; Haar & Donchin, 2020). In fact, the cerebellum is known to modulate activities in different cortical areas during active movements (e.g., the somatosensory cortex, Blakemore et al., 1999). Additionally, the cerebellum’s projection to the posterior parietal cortex (Clower et al., 2001) has been implicated for generating sensory prediction(e.g., Della-Maggiore et al., 2004; Desmurget & Grafton, 2000; also see Blakemore & Sirigu, 2003 for a detailed review).

Is it possible that the cerebellum works in concert with the auditory cortex to compute auditory prediction errors? The cerebellum is certainly known for its involvement in auditory processing (e.g., Aitkin & Boyd, 1975, 1978; Ohyama et al., 2003) including speech perception (Ackermann et al., 2007; Mathiak et al., 2002; Schwartze & Kotz, 2016; Skipper & Lametti, 2021). It is also known that the cerebellum projects to the medial geniculate body (MGB), and the resulting inhibition and/or potentiation of MGB neurons may lead to rapid plasticity of receptive fields of the primary auditory cortex, modulating auditory inputs (e.g., McLachlan & Wilson, 2017; Weinberger, 2011). Such rapid plasticity of the response fields may prepare the primary auditory cortex for discriminating different sounds (David et al., 2012), a function that may be involved in computing auditory prediction errors. Indeed, both right cerebellar areas and bilateral superior temporal cortex were found to be active during speech response to unexpected auditory error (i.e., under the presence of auditory prediction errors, Tourville et al., 2008).

While some studies have suggested that there is no direct projection from the primary auditory area to the cerebellum in primates (e.g., Schmahmann & Pandya, 1991) and mice (e.g., Henschke & Pakan, 2020), others have reported auditory fibers projecting from the superior temporal gyrus and higher-order auditory regions to the cerebellum in primates (e.g., Brodal, 1979). In addition, it is also known that cortical auditory areas project to the cerebellar hemisphere through the cerebro–pontine pathways in some mammals including humans (e.g., Glickstein, 1997; Pastor et al., 2008). Collectively, while exactly how neurons in auditory regions compute auditory prediction errors remain unclear, it is certainly likely that they are estimated through several pathways incorporating multiple cortical and cerebellar areas.

It is also noteworthy that baseline SIS activities are found to be most pronounced in the left auditory cortex, in line with the notion that the left hemisphere is dominant in speech and language perception (Curio et al., 2000; Houde et al., 2002). In this study, we found SIS reduction in the left auditory cortex alone, in line with a previous study that found prediction-related SIS effect only in the left hemisphere (Niziolek et al., 2013). One discrepancy between the current study and Niziolek et al. (2013) is that we did not find a significant SIS effect in the right hemisphere even during the baseline phase (see Supplementary Information 2). Given that the right hemisphere SIS is known to be highly variable across tasks and individuals (K. X. Kim et al., 2023), the discrepancy may have been due to the sampling issue.

### Limitations and future directions

The current study examined speaking-induced suppression during auditory-motor adaptation and found evidence consistent with the notion that sensory prediction errors drive implicit adaptation. However, some limitations must also be noted. First, we excluded two participants who showed “following” behaviors given that the scope of the paper was to examine SIS during adaptation (see Methods for more details). Nonetheless, a recent study points out that such behavior may represent the tail of a unimodal distribution across the participants (Miller et al., 2023). In light of this new finding, future studies should examine adaptation-related SIS reduction in a large number of participants for higher statistical power, rather than excluding “following” responses. Another limitation of the study is that we repeated adaptation sessions to acquire an adequate number of trials for SIS analyses, which introduces noise in the data given that adaptation behaviors may differ in the two sessions (see Supporting Information 1). With continual advancements in developing powerful source reconstruction algorithms, it may be possible to acquire enough trials from a single adaptation session in future SIS adaptation studies.

Additionally, previous studies in monkeys (Eliades & Wang, 2017) and humans (Chang et al., 2013) reported that different neurons in the auditory areas behave differently during speaking (or vocalization) vs. listening conditions. Future investigations on which types of these neurons in the auditory cortex are ultimately responsible for SIS reduction during adaptation will be critical for understanding how sensory prediction errors are generated in the central nervous system. Lastly, future studies should examine these questions in more ecologically valid contexts given the recent finding of SIS (Kurteff et al., 2023) and adaptation (Lametti et al., 2018) in more naturalistic tasks such as sentences.

## Methods

### Participants

Participants who were native speakers of American English with no known communication, neurological, or psychological disorders were recruited for both MEG and MRI recording sessions. 30 participants participated in the adaptation experiment, but 8 participants were excluded from analyses for various reasons. One participant’s source could not be reliably localized, and four participants could not finish the task due to fatigue. Two participants showed “following” non-adaptive behavior (i.e., change of 15 Hz or more in the direction of the perturbation in the late learning phase) and one participant had atypical (outlier) SIS response in the baseline, (SIS < –10 z).

Here, we report adaptation experiment results from the remaining 22 participants (mean age = 30.9, SD = 8.8 years old, 12 females). For the control experiment, 12 participants (mean age = 31.6, SD = 8.0 years old, 5 females) participated. It should be noted that 5 of these participants also participated in the adaptation experiment, each visit being 1-2 months apart. The order was pseudorandomized (i.e., three of the five participants did the adaptation experiment first). All participants in the study passed pure-tone hearing thresholds of ≤ 20 dB HL for the octave frequencies between 500 and 4,000 Hz, except one participant in the adaptation experiment whose threshold was at 30 dB in the right ear and 40 dB in the left ear at 4 kHz.

### Tasks

#### Adaptation experiment

During MEG data collection of the first two sessions, participants were asked to read “Ed,” “end,” or “ebb” (60 trial blocks for 3 different words = 180 total trials) that appeared on the screen. During these speaking sessions, participants heard their speech with the first formant frequency (Formant 1 or F1) shifted upward for some trials (trial blocks 21 to 50, see below), which made their speech to sound more like “Add,” “And,” and “Ab,” respectively. The auditory perturbation, 150 Hz upshift, was applied through Feedback Utility for Speech Processing (FUSP, Kothare et al., 2020) and the total feedback latency (i.e., hardware + software, K. S. Kim et al., 2020) was estimated to be about 19 ms.

During the speaking sessions, the first 20 trial blocks (i.e., baseline) had no perturbation, while blocks 21 through 50 had a 150 Hz up-shift perturbation in the auditory feedback. We categorized the first 15 trial blocks of the perturbed trials (21 – 45) as the early learning phase and the second 15 trial blocks (36-50) as the late learning phase. In the passive listening condition, participants heard the same auditory feedback that they received during the speaking condition (including the perturbed sounds) through the earphones. With a mean interstimulus interval of 3s and short breaks (roughly 20 seconds) every 30 utterances, the duration of each session was approximately 10 – 12 minutes.

Participants performed two speaking sessions, after which they performed two listening sessions. Between each session, participants were given a few minute-long break that included conversations with the experimenter, which allowed additional exposure to their unaltered auditory feedback. Given that the adaptation task (i.e., speaking session) was repeated, we also checked whether there was any savings effect and found that there was no consistent effect of repeating adaptation (see Supplementary Information 1).

#### Control experiment

We also designed a control experiment in which we applied 0 Hz perturbation (instead of 150 Hz perturbation) during early and late “learning” phases. All other details of the task remained identical to the adaptation experiment.

#### MRI

On a separate day, participants also underwent an MRI scan, where a high-resolution T1-weighted anatomical MRI was acquired in each participant for source reconstruction.

#### MEG acquisition

Participants were placed in a 275-channel, whole-head biomagnetometer system (Omega 2000, CTF, Coquitlam, BC, Canada; sampling rate 1200 Hz; acquisition filtering 0.001-300 Hz) for a total of four sessions (two speaking and two listening sessions). Participants heard auditory feedback (or recorded auditory feedback during listening condition) through ER-3A ear-insert earphones (Etymotic Research, Inc., Elk Grove Village, IL) and a passive fiber optic microphone (Phone-Or Ltd., Or-Yehuda, Israel) was placed about an inch in front of their mouths to record speech responses. All stimulus and response events were integrated in real time with MEG timeseries via analog-to-digital input to the imaging acquisition software.

Each participant lay supine with their head supported inside the helmet along the center of the sensor array. Three localizer coils affixed to the nasion, left peri-auricular, and right peri-auricular points determined head positioning relative to the sensor array both before and after each block of trials. We ensured that participants’ head movements were smaller than 5 mm in every session. Co-registration of MEG data to each individual’s MRI image was performed using the CTF software suite (MISL Ltd., Coquitlam, BC, Canada; ctfmeg.com; version 5.2.1) by aligning the localizer coil locations to the corresponding fiducial points on the individual’s MRI. MRI images were exported to Analyze format and spatially normalized to the standard T1 Montreal Neurological Institute (MNI) template via Statistical Parametric Mapping (SPM8, Wellcome Trust Centre for Neuroimaging, London, UK).

### Data extraction and analyses

#### First formant frequency (F1)

The first formant frequency (F1) from each speech production was extracted through a custom MATLAB software, Wave Viewer (Raharjo et al., 2021). We then extracted F1 from the vowel midpoint (40% to 60% into the vowel) and averaged it for each utterance. In case of missing trials, we replaced the data point by using an interpolation method using four nearest neighboring trials as described in Kitchen et al. (2022). We replaced about 2.96% and 2.88% of the data for the adaptation and control experiments respectively. We normalized the data by subtracting the baseline F1 from the data (i.e., baseline = 6^th^ to 20^th^ trial blocks). The amount of learning in each phase was assessed by averaging the last 5 trial blocks (31^st^ to 35^th^ blocks for early learning and 46^th^ to 50^th^ blocks for late learning).

#### Speaking-induced suppression

We first corrected distant magnetic field disturbances by calculating a synthetic third-order gradiometer, detrended using a DC offset across whole trials, and then filtered (4th order Butterworth, bandpass 4 to 40 Hz) sensor data. In the sensor data of two participants (one of them participated in both of the experiments, so three datasets), considerable (>10pT) sensor noise caused by dental artifact verified through visual inspection was denoised using a dual signal subspace projection (DSSP, Cai, Kang, et al., 2019; Cai, Xu, et al., 2019). After pre-processing sensor data, separate datasets were created with trials during baseline, early learning, and late learning phases for speak and listen conditions. In these datasets, trials exceeding a 2 pT threshold at any timepoint were rejected. In two participants’ data, three channels were removed prior to threshold-based artifact rejection. The data was then averaged across all remaining channels. For the adaptation experiment, 5.47% of the speak session trials and 4.70% of the listen session trials were removed. For the control experiment, 5.68% and 6.14% of the trials were removed for speak and listen sessions, respectively.

For each participant, a single-sphere head model was derived from the individual’s co-registered T1 structural MRI using the CTF software suite (MISL Ltd., Coquitlam, BC, Canada; ctfmeg.com; version 5.2.1). Using the Champagne algorithm (Owen et al., 2012) and a lead field of 8mm resolution on the baseline listen data, we generated whole-brain evoked activity between 75 ms and 150 ms (after the auditory feedback onset), and determined the MNI coordinate with the most pronounced M100 response in the left and right auditory areas (i.e., the highest amplitude) for each participant. Although we only report the results from the left auditory area in the main text, the results for the right hemisphere can be found in Supplementary Information 2. The median MNI coordinate across adaptation and control experiments were [x = –56, y = –24, z = 8] and [x = 48, y = –24, z = 8] for the left and right auditory areas respectively. We then used a Bayesian adaptive beamformer (Cai et al., 2023) to extract time-series source activity focused on the obtained MNI coordinate across all phases (i.e., baseline, early, and late). From the final time-series z-scored data, we measured M100 peak by finding the maximum value between 75 – 150 ms after the auditory signal. We then computed the M100 amplitude difference between the listen and speak sessions to determine SIS:

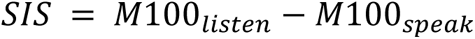

#### Statistical analysis

A linear mixed effects model was constructed for SIS with the different adaptation phases as fixed effects and participants as a random effect using *lme4* package in R (Bates et al., 2015). The Tukey test was used for post-hoc pairwise comparisons from the *emmeans* package in R (Lenth, 2022). A Pearson’s correlation tested to examine relationships between the amount of adaptation and the SIS amplitudes.

## Supplementary Information

(Supplementary Information 1, 2, 3 can be found in a separate document)

## Acknowledgement

This study was funded by grants from the National Institute of Health (F32DC019538 to K.S.K., R01DC017696, R01DC017091, R01DC013979, R01NS100440, and P50DC019900 to J.F.H., S.S.N.). The funders had no role in study design, data collection and analysis, decision to publish, or preparation of the manuscript. We would also like to thank Ashley Tay, Joshua Chon, Mahmoud Jiha, Derek Kinsella, Gavin Belok, and Lingwei Ouyang for their help with data collection and extraction.

