## Supplementary Information for "Neurophysiological evidence of sensory prediction errors driving speech sensorimotor adaptation"

**Supplemental Information 1: Repeated adaptation session**

Unlike arm reaching adaptation studies showing that re-experiencing adaptation results in savings (e.g., Huang et al., 2011) or attenuation in case of implicit adaptation (Avraham et al., 2021), we did not find any significant changes on the repeated adaptation session (2^nd^ session, see Fig. S1A). The un-normalized baseline phase (before normalizing the baseline phase to 0 Hz) showed no significant difference across the two sessions, t(21) = –2.344, p_adj_ = 0.087 (Fig. S1B, left), though the p-value was close to α = 0.05 due to a few participants whose baseline F1 was much lower in the 2^nd^ session than the 1^st^ session. Nonetheless, there was no clear trend on whether these individuals learned more or less in the 2^nd^ session. Indeed, adaptation across the repeated adaptation sessions was not different in both early learning, t(21) = –0.830, p_adj_ = 0.416, and late learning, t(21) = 1.062, p_adj_ = 0.300, (see Fig. S1B, right).

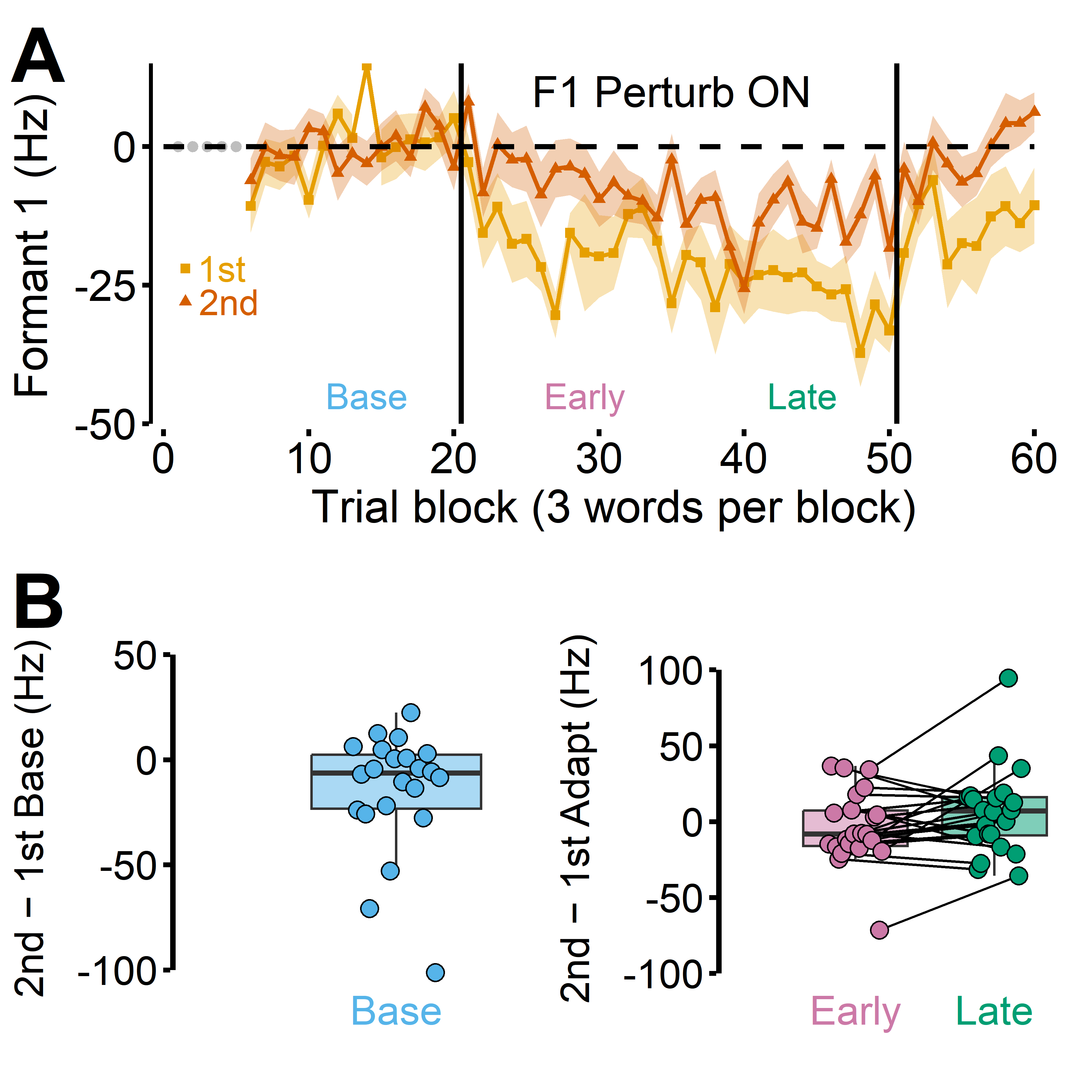

Fig. S1. A: Adaptation did not differ between the first and second sessions. B: The overall baseline also did not change in most participants (left). There were two participants whose baseline in the second session was reduced by more than 60 Hz, but as a group the baseline did not differ between the two sessions, t(21) = –2.344, p_adj_ = 0.087. Overall, participants as a group, there was no sign of savings or attenuation in the second session (right).

**Supplemental Information 2: Right hemisphere data**

Compared to the left auditory cortex, the right hemisphere SIS activities were less pronounced. Multiple individuals did not show a clear SIS response in the right hemisphere even during the baseline in the adaptation group, t(21) = 1.148, p = 0.264 (see Fig. S2, left). In the control group, however, SIS was significant, t(11) = 2.780, p = 0.018 (see Fig. S2, right). In the adaptation group, we did not observe a significant SIS reduction in adaptation phases in the right auditory cortex, F (2, 44) = 0.934, p = 0.401, in line with a previous study that found prediction-related SIS effect only in the left hemisphere (Niziolek et al., 2013). We also did not find any significant SIS reduction in the control group’s right hemisphere activities, F (2, 24) = 0.469, p = 0.631.

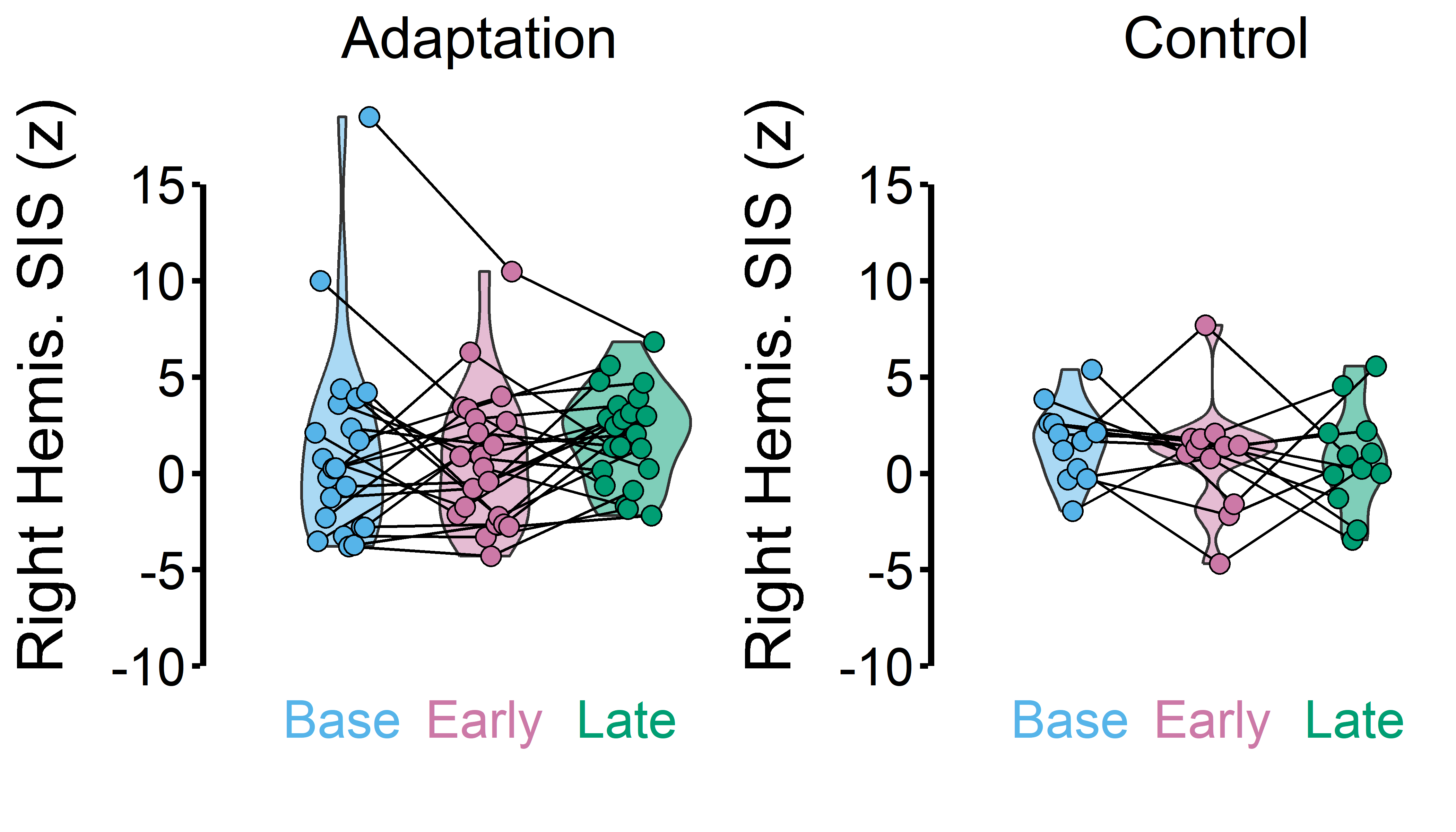

Fig. S2. We did not find a significant SIS effect during the baseline in the right hemisphere in the adaptation group, t(21) = 1.148, p = 0.264. We did not observe any significant SIS reduction during adaptation in both the adaptation and control groups.

**Supplemental Information 3: Late SIS reduction (from baseline SIS) vs. Final adaptation**

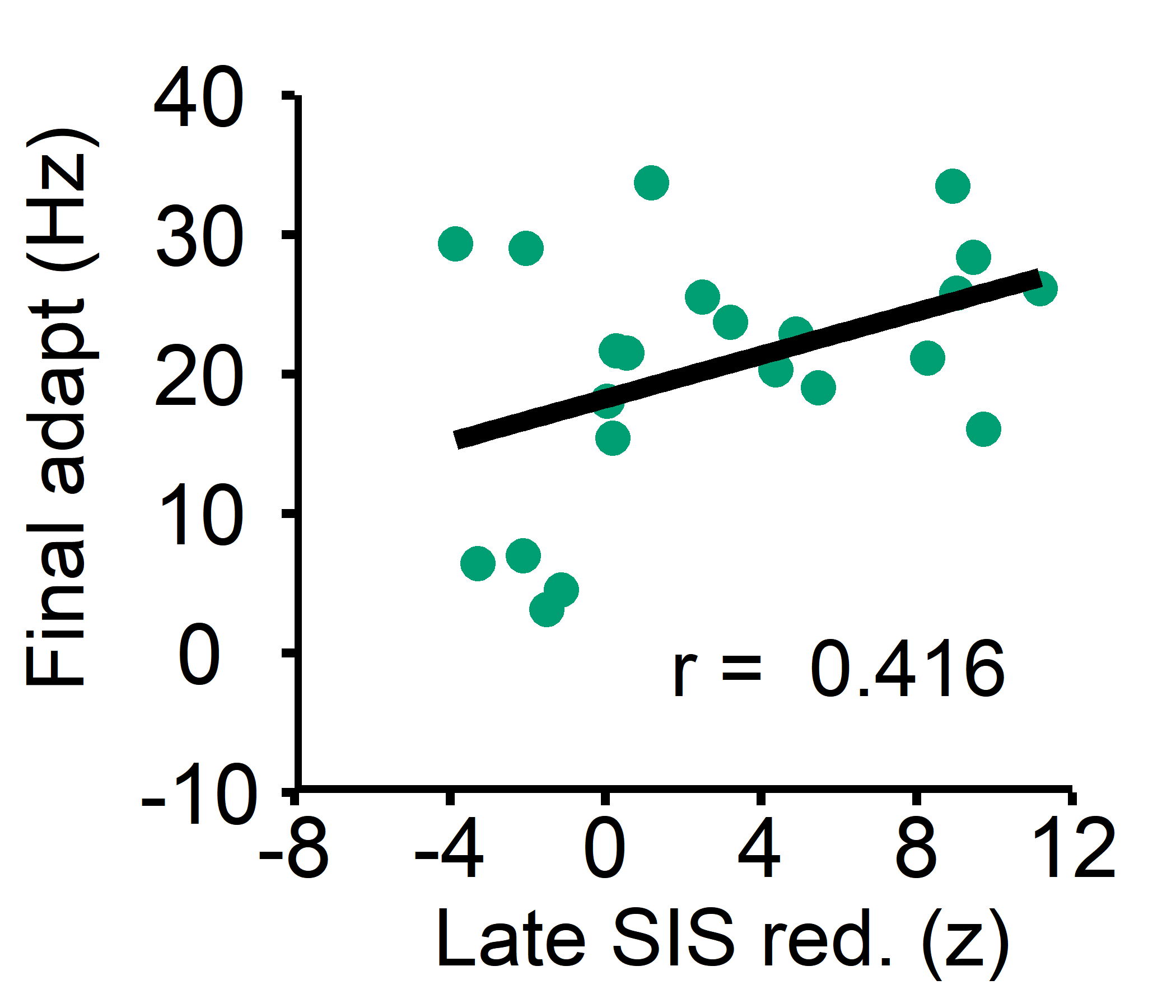

Fig. S3. We examined whether late SIS reduction from the baseline SIS could predict the final amount of adaptation. They were not significantly correlated, though trending, r(20) = 0.416, p = 0.054.
